# Standalone nanopore sequencing for foodborne pathogen surveillance: a large-scale evaluation and quality control framework

**DOI:** 10.64898/2026.03.20.713089

**Authors:** Michael Biggel, Nicole Cernela, Jule Horlbog, Michael S. DeMott, Peter C. Dedon, Michael B. Hall, Jessica Chen, Peyton Smith, Heather A. Carleton, Roger Stephan, Lara Urban

**Affiliations:** Institute for Food Safety and Hygiene, University of Zurich, Zurich, Switzerland; National Reference Centre for Enteropathogenic Bacteria and Listeria (NENT), Zurich, Switzerland; Department of Biological Engineering, Massachusetts Institute of Technology, Cambridge, MA, USA; The University of Queensland, UQ Centre for Clinical Research, Herston, QLD 4029, Australia; Enteric Diseases Laboratory Branch, Centers for Disease Control and Prevention, Atlanta, GA, USA

**Author notes:** **Correspondence**: Michael Biggel, Winterthurerstr. 272, 8057 Zurich.

**Keywords:** Oxford Nanopore Technologies, Whole-genome sequencing, Foodborne pathogen surveillance, Outbreak detection

## Abstract

Whole-genome sequencing (WGS) is central to foodborne pathogen surveillance and cross-border outbreak detection. Long-read sequencing using Oxford Nanopore Technologies (ONT) promises rapid, complete, and cost-effective genome assemblies in a single workflow. However, the adoption of standalone ONT sequencing of native DNA has been slowed by concerns that DNA modifications can compromise per-base sequencing accuracy and downstream genotyping.

In this study, we evaluated ONT-only sequencing performance across 294 genetically diverse isolates representing ten major foodborne pathogens. Using the SUP@v5.2 basecalling model at 50x coverage, 97.3% (286/294) of the ONT assemblies produced identical or near-identical cgMLST profiles (≤3 allelic differences) as Illumina-polished hybrid assemblies. Elevated error rates were observed in four *Salmonella enterica* serovar Kentucky and four *Listeria monocytogenes* isolates and were associated with the presence of specific DNA phosphorothioation or methylation systems.

Re-basecalling the same dataset with the newly released HAC@v6.0 model revealed a different error profile: although 93.5% (275/294) of assemblies remained highly accurate, all 13 isolates carrying *dnd* (DNA phosphorothioation) or *dpd* (7-deazaguanine modification) systems, including isolates of *S. enterica*, *Cronobacter sakazakii*, and *Vibrio parahaemolyticus*, exhibited high error rates, suggesting that such atypical modifications were not adequately represented in the model’s training dataset.

To enable rapid identification of unreliable assemblies, we developed alpaqa, a lightweight computational tool that detects systematic nanopore assembly errors without requiring supplemental short-read data or reference genomes. By identifying affected assemblies, alpaqa provides a quality safeguard for ONT-only workflows. Masking low-quality bases in assemblies flagged by alpaqa improved cgMLST accuracy, although this reduced the number of callable loci and therefore genotyping resolution.

Our findings demonstrate that standalone ONT sequencing of native DNA is sufficiently accurate for routine foodborne pathogen surveillance when combined with appropriate quality control, supporting its use in harmonised genomic surveillance frameworks.

**Data summary:** All sequencing data generated in this study have been submitted to the NCBI Sequence Read Archive. Accession numbers for Illumina and ONT (SUP@v5.2) reads are listed in **Supplementary Table S1**. Raw pod5 files from error-prone isolates have been deposited in SquiDBase (SQB000021). Alpaqa is available at github.com/MBiggel/alpaqa/. An automated ONT assembly and quality control pipeline integrating alpaqa is available at github.com/MBiggel/boap/.

**Impact statement:** This study demonstrates that standalone Oxford Nanopore sequencing of native DNA can achieve highly accurate genotyping for routine foodborne pathogen surveillance across diverse species. We show that the remaining inaccuracies are linked to specific DNA modification systems, including phosphorothioation and 7-deazaguanine modifications, which are identified here as previously unrecognised sources of systematic sequencing errors. To address this limitation, we introduce alpaqa, a reference-free method for detecting such error-prone assemblies, providing a practical quality-control framework for ONT-only workflows. Together, these results support the reliable use of nanopore sequencing in routine genomic surveillance.

## Introduction

Whole-genome sequencing (WGS) has revolutionized our ability to understand, track, and combat bacterial pathogens. By enabling the discrimination between closely related strains, reconstruction of transmission routes, and detection of antimicrobial resistance and virulence determinants, WGS has become central to modern pathogen surveillance and cross-border outbreak detection [1, 2]. The integration of WGS into routine surveillance is a strategic priority for public health agencies worldwide to strengthen genomic surveillance and outbreak response [3–5]. For over a decade, Illumina short-read sequencing has been the gold standard for WGS for pathogen surveillance due to its high per-base accuracy and has enabled harmonised genomic typing approaches across public health laboratories, facilitating international data sharing and coordinated outbreak investigations. However, short reads frequently fail to resolve repetitive regions and structural variants, resulting in fragmented assemblies [6].

Long-read nanopore sequencing by Oxford Nanopore Technologies (ONT) overcomes this limitation by enabling the complete assembly of bacterial chromosomes and plasmids. In addition to improved genome reconstruction, ONT offers an accessible and cost-effective infrastructure, rapid turnaround times, and scalable throughput, making it an attractive platform for pathogen surveillance [7]. Early ONT versions suffered from high error rates that required polishing with short-read data to achieve reliable assembly accuracy [8]. However, recent advances in flow cell technology (R10.4), chemistry, basecalling, and polishing algorithms have substantially enhanced raw read and consensus accuracy. High-quality bacterial genomes can now be generated from ONT data alone, with accuracy levels for core genome multi-locus sequence typing (cgMLST) and variant calling approaching – and in repetitive regions sometimes exceeding – Illumina-based approaches [9–14]. As a result, nanopore sequencing is transitioning from a complementary technology to a standalone platform for genomic surveillance, offering advantages for both clinical and food safety surveillance where rapid and reproducible genomic typing is essential for outbreak detection and source attribution [15, 16].

Despite technological improvements, concerns remain regarding the potential impact of epigenetic DNA modifications, such as methylations, on ONT basecalling accuracy. Although improved basecalling and polishing algorithms account for common modifications, rare or lineage-specific modification systems may still introduce systematic assembly errors that could lead to the incorrect exclusion of epidemiologically related isolates from outbreak clusters [17–19]. The prevalence and impact of such systems among diverse foodborne pathogens remain insufficiently characterised. Comprehensive evaluations across large and genetically diverse isolate collections are needed to determine the reliability and limitations of ONT-only WGS for routine genomic surveillance workflows.

In this study, we systematically evaluate the accuracy of standalone ONT-based WGS across 294 isolates representing diverse lineages of ten major foodborne bacterial pathogens commonly targeted in public health surveillance. Using read downsampling, we assess the impact of sequencing depth on assembly accuracy and compare the current high-accuracy (HAC) and super-accuracy (SUP) dorado basecalling models, which differ in their trade-off between computational efficiency and basecalling accuracy. We identify atypical DNA modification systems associated with systematic sequencing errors, highlighting gaps in current basecaller training datasets. To address this challenge, we introduce alpaqa, a computational approach that detects unreliable assemblies directly from consensus quality scores without requiring reference genomes or short-read validation, supporting the use of ONT sequencing in routine foodborne pathogen surveillance.

## Results

### High cgMLST accuracy of ONT-only assemblies

We generated Illumina and ONT (RBK114, SUP@v5.2 basecalling) WGS data for 294 isolates representing major foodborne pathogens, with minimum sequencing depths of 40x (Illumina) and 50x (ONT) coverage. Isolates belonged to the *Bacillus cereus* group (n = 19), *Campylobacter* (n = 62), *Cronobacter sakazakii* (n = 15), *Listeria monocytogenes* (n = 78), *Salmonella enterica* (n = 57), *Shigella* (n = 16), *Vibrio parahaemolyticus* (n = 8), and *Yersinia enterocolitica* (n = 39).

At a downsampled depth of 50x, ONT-only assemblies were highly accurate. Overall, 97.3% (286/294) of the isolates yielded identical or near-identical (≤3 allelic differences) cgMLST profiles compared to their corresponding Illumina-polished reference assemblies (**Figure 1**), including 86.7% (255/294) with fully identical profiles. However, we observed elevated error rates in four *L. monocytogenes* and four *S. enterica* assemblies, which correlated with the presence of specific DNA modification systems.

**Figure 1.**
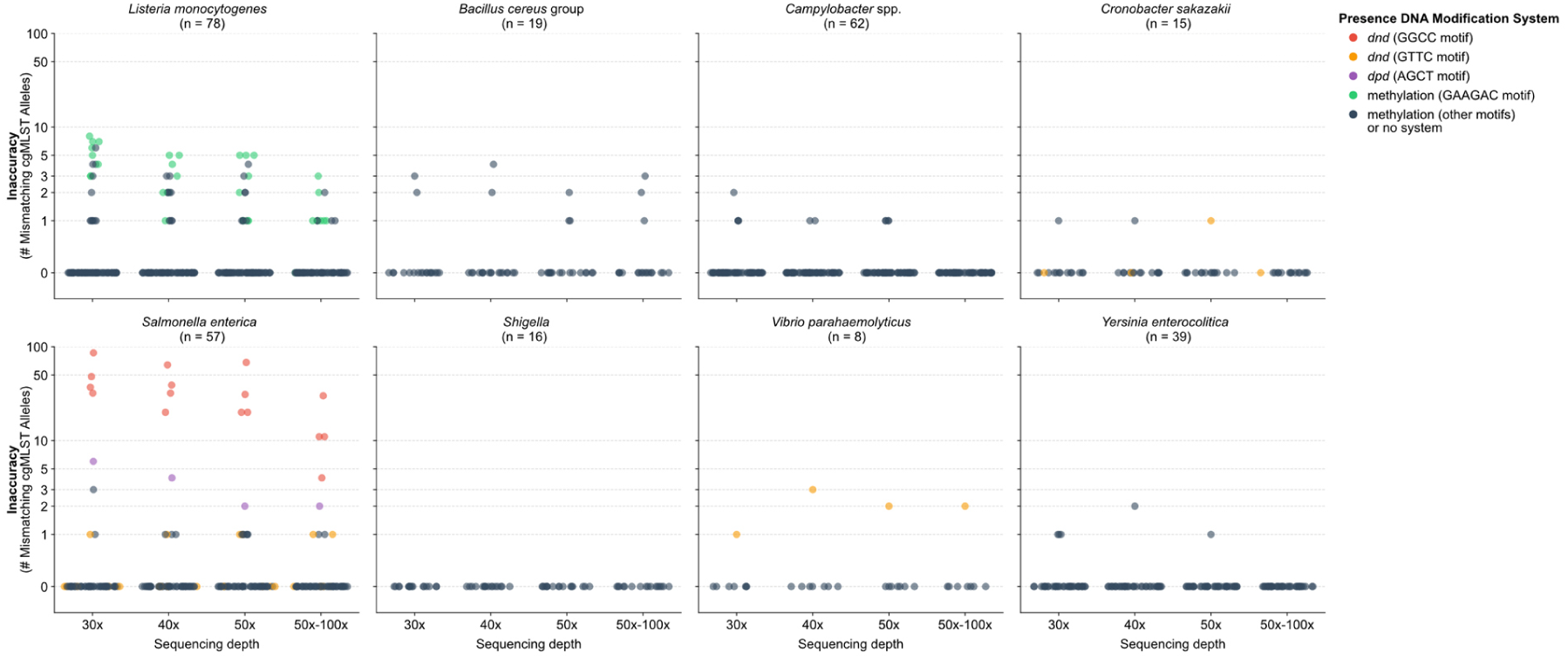
Accuracy of 294 ONT-only assemblies from SUP@v5.2 data at varying sequencing depths. The number of mismatching cgMLST alleles was determined by comparing ONT-only assemblies generated at 30x, 40x, 50x, and full depth (up to 100x) to their corresponding hybrid assembly. Each dot represents the assembly of a single isolate, with colors indicating the presence of different modification systems. The y-axis is plotted on a logarithmic scale.

### DNA modification systems associated with elevated assembly error rates

The four error-prone *L. monocytogenes* isolates exhibited four to five mismatching alleles at 50x depth (**Figure 1**). These errors were previously attributed to methylation by specific Type II restriction-modification systems targeting GAAGAC (5′-GAAG6mAC-3′/5′-GT4mCTTC-3′) or GCNGC (G5mCNGC) [18]. Increased sequencing depth of 100x reduced the number of mismatching cgMLST alleles to ≤3 mismatches.

The four error-prone *S. enterica* isolates all belonged to serovar Kentucky (ST152, ST314, or ST696) and showed substantially higher error rates, with 20 to 68 mismatching alleles (**Figure 1**). Increasing sequencing depth or an alternative assembly approach using autocycler followed by dorado polishing reduced the number of mismatching cgMLST alleles (to 4-30 and 6-36, respectively), but neither approach resolved the inaccuracies. Errors consistently occurred at GGCC sites. While no shared exclusive methyltransferases were identified among these isolates (**Supplementary Table S2**), all carried a specific *dndACDE* phosphorothioation system (**Supplement S1**). LC-MS/MS analysis confirmed phosphorothioate-linked GPSG dinucleotides in these isolates, supporting GGCC as the target recognition site of this *dnd* system. Although the operon lacked the repressor *dndB*, GPSG was observed at average frequencies of 6 to 7 modifications per 10^4^ nucleotides, which is comparable to levels detected in isolates with complete *dnd* systems [20]. Screening of public genomes showed that the Kentucky-associated *dnd* system is present in all assemblies belonging to the ST152/ST314-associated *S. enterica* serovar Kentucky clade (**Supplement S2**). In public repositories, it was also detected in two assemblies belonging to *S. enterica* serovar 50:z:e,n,x NCTC9936 (GCA_900478195.1) and *K. variicola* KLE2 (GCA_050990805.1) (**Supplement S1**). In contrast, accurate assemblies were obtained for eight *C. sakazakii*, *S. enterica*, and *V. parahaemolyticus* isolates harbouring the reference *dnd* system (*dndBCDE*) originally described in *S. enterica* serovar Cerro 87 with the target motif 5′-GPSAAC-3′/5′-GPSTTC-3′ (**Figure 1**) [21, 22].

One additional *S. enterica* isolate (serovar Montevideo, N21-3953) exhibited slightly elevated error rates (two mismatching alleles at 50x coverage) and carried a *dpd* DNA modification system responsible for 7-deazaguanine modifications [23].

### Assembly accuracy remains high at reduced sequencing depth and with HAC@v5.2 basecalling

To estimate the impact of lower sequencing depths on assembly accuracies, we downsampled ONT reads to 40x and 30x coverage. Most assemblies (280/294, 95.2%) remained highly accurate (≤3 mismatching alleles) even at 30x coverage (**Figure 1**). Basecalling with the high-accuracy HAC@v5.2 model also produced largely reliable assemblies, with 92.2 % (271/294) maintaining ≤3 mismatching alleles at 50x depth, although overall accuracy was lower than with the SUP model.

### High error rates in *dnd*+ and *dpd*+ assemblies from HAC@v6.0 basecalled data

Basecalling with the HAC@v6.0 model resulted in 93.5 % (275/294) assemblies with ≤3 mismatching alleles at 50x depth. Whereas assemblies from isolates with methylation systems or no detected modification systems had accuracies comparable to those generated with SUP@v5.2, all 13 assemblies with *dnd* or *dpd* modification systems consistently exhibited high error rates (24 to 314 mismatching alleles, median 78) (**Supplementary Table S1**). To assess the broader relevance of these findings, we screened publicly available genome assemblies from 67 clinically important bacterial pathogens. We detected *dnd* systems in 44 of 67 bacterial species, with prevalence rates of more than 10% in several species such as *Mycobacterium abscessus*, *Citrobacter koseri*, and *Clostridioides difficile* (**Supplement S3**). *dpd* systems were largely restricted to Enterobacterales, with prevalence rates of up to 8% in several *Klebsiella*, *Escherichia*, and *Enterobacter* species.

### Detection of unreliable assemblies using the alpaqa quality assessment tool

Our analysis confirms that ONT sequencing with SUP basecalling achieves a high accuracy for >95% of the investigated isolates at commonly used sequencing depths. However, the remaining inaccurate assemblies represent a challenge for routine surveillance. Currently, detecting such unreliable assemblies requires Illumina data for cross-validation, limiting ONT’s use as a standalone method. To address this, we developed alpaqa, a lightweight bioinformatic tool designed to flag error-prone assemblies using ONT data alone. Alpaqa is based on the observation that DNA modifications not recognized by the basecalling model typically produce very low-quality scores in the consensus assembly, often dropping below Q5 (**Figure 2**). Alpaqa analyses per-base quality scores generated during assembly polishing and evaluates the distribution of low-quality bases (LQBs) across the longest contig. By first masking LQB-dense regions linked to artefacts and then calculating a normalized LQB density per Megabase (Mbp), alpaqa provides a quantitative metric for assembly reliability. In addition, alpaqa identifies k-mers that are significantly enriched for LQBs, enabling detection of error-associated motifs such as GGCC and GAAGAC.

In our dataset, LQB density correlated with cgMLST accuracy (**Figure 3A**; **Supplementary Table S3**). At 50x depth, 285 of 286 (99.7%) assemblies with fewer than 5 LQBs/Mbp were confirmed as highly accurate (≤3 allelic mismatches). The single outlier was an error-prone *L. monocytogenes* assembly with five mismatching alleles and 4.5 LQBs/Mbp. Assemblies exceeding this threshold were consistently inaccurate. The three remaining error-prone *L. monocytogenes* showed 6 to 8 LQBs/Mbp, whereas the error-prone *S. enterica* Kentucky assemblies exhibited substantially higher LQB densities (14 to 37 LQBs/Mbp). The correlation between LQB density and assembly accuracy remained robust for the HAC@v5.2 and HAC@v6.0 datasets, although the predictive separation was less distinct (**Figure 3B**). Here, several accurate assemblies showed LQB densities between 5 and 10 LQBs/Mbp. Of the 13 error-prone *dnd*+/*dpd*+ assemblies generated with HAC@v6.0, 12 exhibited >10 LQBs/Mbp, while the remaining assembly had 9.25 LQBs/Mbp (**Figure 3C**). Across both SUP and HAC datasets, assemblies with fewer than 2.5 LQBs/Mbp were consistently highly accurate.

**Figure 2.**
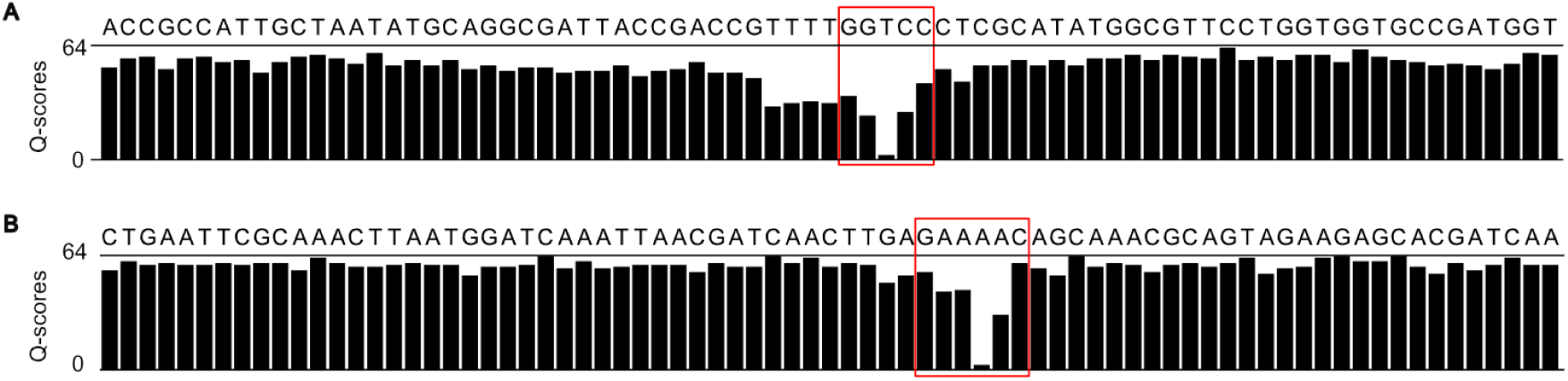
Per-base consensus quality scores across representative genomic assembly windows. Bar plots show the Phred-scaled quality scores for each base in the final assembly. (A) *Salmonella enterica* Kentucky N18-0332, carrying a putative GGCC-targeting phosphorothioation system. (B) *Listeria monocytogenes* N19-1094, containing a GAAGAC-targeting methyltransferase. Red squares indicate the incorrectly called recognition motifs of the DNA modification systems, each containing a low-quality base (LQB). The genome-wide density of LQBs can be used as a predictor of systematic errors and overall assembly accuracy.

**Figure 3.**
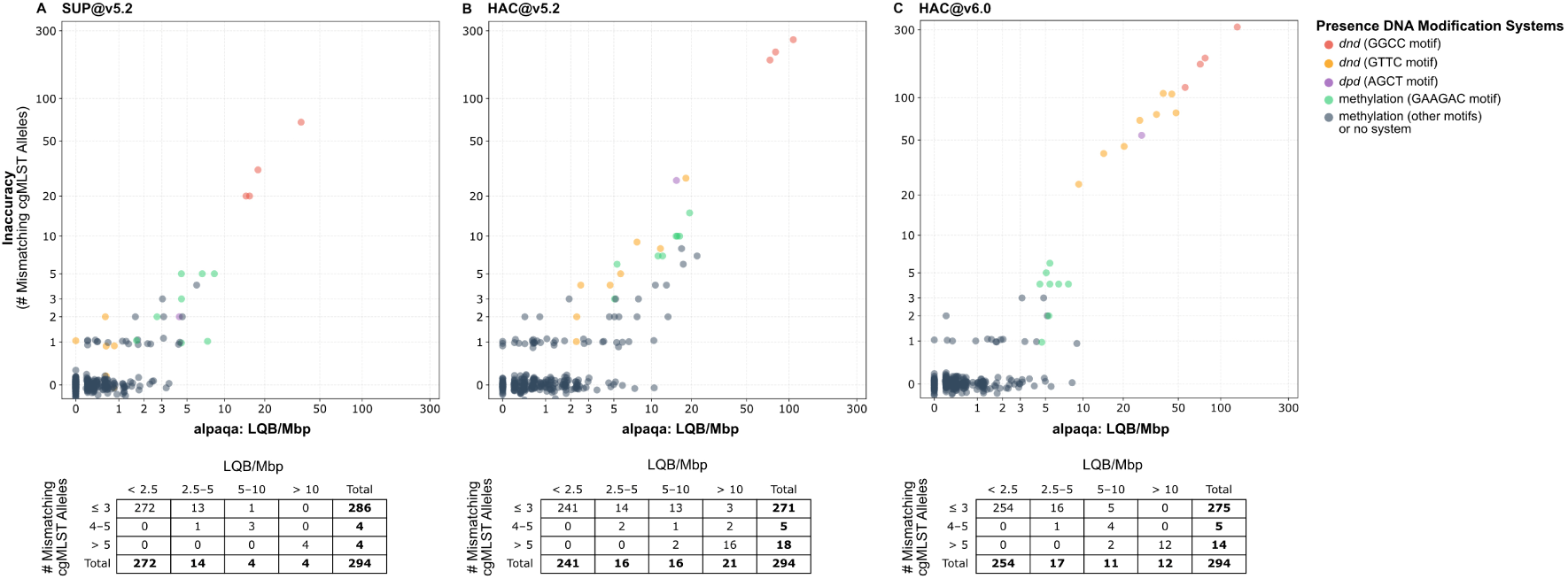
Correlation between LQB density and cgMLST allelic mismatches. The scatterplots illustrate the relationship between the frequency of low-quality bases (LQB) per Mbp determined with alpaqa and the number of mismatching cgMLST alleles across 294 ONT-only assemblies (50x target coverage) from data basecalled with (A) SUP@v5.2, (B) HAC@v5.2, and (C) HAC@v6.0. Each dot represents an assembly, with colors indicating the presence of specific modification systems. A small vertical jitter was applied to dots with 0 and 1 mismatching alleles to reduce overlap. Both axes are plotted on a logarithmic scale. Tables summarize the frequency of assemblies across different LQB densities and allelic mismatch counts.

To further validate the generalisability of the alpaqa metric, we applied the tool to publicly available data from 147 previously benchmarked isolates from multiple clinically relevant bacterial species, confirming that low LQB density is a strong predictor of high cgMLST accuracy (**Supplement S4, Supplementary Table S4**).

### Masking of low-quality bases improves cgMLST accuracy of error-prone assemblies

To evaluate whether masking low-quality bases improves genotyping accuracy, assemblies were processed using fastq2a prior to cgMLST analyses. Because masked bases ("N") are excluded from allele calling in pyMLST or Ridom SeqSphere, this approach converts potential sequencing errors into missing data.

For the four inaccurate *L. monocytogenes* assemblies, masking bases with Q≤10 excluded 18 to 27 error-associated loci (∼1.5 % of 1701 cgMLST targets) and reduced mismatches to ≤1 allele. For the four inaccurate *S. enterica* assemblies, masking bases with Q≤10 excluded 82 to 205 error-associated loci (∼3 to 7 % of 3002 targets) and reduced mismatches to 4 to 12 alleles. Here, achieving near-accurate profiles (0 to 4 mismatches) required a stricter threshold (Q≤15), but this excluded substantially more loci (178 [5.9%] to 372 [12.4%]), reducing cgMLST resolution (**Supplement S5**).

Our fully automated ONT assembly pipeline, BOAP, integrates alpaqa-based quality assessments and the masking of low-quality bases in unreliable assemblies to enable standardised ONT-based genomic surveillance.

## Discussion

Current ONT sequencing workflows enable highly accurate WGS for pathogen surveillance. Across 294 genetically diverse bacterial isolates representing ten major foodborne pathogens, 97.3% of the ONT-only assemblies from SUP@v5.2 data produced identical or near-identical cgMLST profiles compared to Illumina-polished references. While these results were achieved at a moderate sequencing depth of 50x coverage, most assemblies (95.2%) remained highly accurate even at 30x coverage. Increasing sequencing depth beyond 50x provided only marginal improvements. These findings demonstrate that ONT sequencing is sufficiently robust for routine cgMLST-based surveillance without requiring excessive sequencing depth.

Despite the overall high accuracy, systematic errors persisted in a small subset of SUP@v5.2 assemblies and were associated with atypical epigenetic modifications. At 50x coverage, affected assemblies showed up to 68 discrepant cgMLST alleles, compromising outbreak and transmission inference. Importantly, such error-prone assemblies are not readily identifiable using standard assembly metrics and may therefore remain undetected in routine ONT-only workflows. To address this limitation, we developed alpaqa, a computational tool that identifies unreliable assemblies directly from ONT data based on consensus quality score distributions. This approach enables reference-free detection of problematic assemblies without requiring orthogonal Illumina validation, removing a major barrier to the adoption of ONT sequencing as a standalone platform for genomic surveillance, including for previously understudied pathogens.

Several strategies can be applied once unreliable assemblies are identified. Masking low-quality bases in assemblies results in their exclusion during cgMLST analysis, substantially improving cgMLST accuracy. However, this approach reduces the number of callable loci and thereby decreases genotyping resolution, which may limit discriminatory power for outbreak investigations, particularly in highly error-prone assemblies. Alternatively, affected isolates may be re-sequenced using Illumina or PCR-based ONT library preparation, which removes native DNA modifications and restores high assembly accuracy [24]. Given the low frequency of problematic isolates (<3%) observed across a diverse pathogen collection, routine amplification-based sequencing is generally not desirable, as it reduces read length, assembly contiguity, and overall workflow efficiency [24]. Instead, selective application of targeted mitigation strategies guided by quality assessment tools may represent a more practical solution. We implemented the fully automated Nextflow pipeline BOAP integrating assembly, polishing, alpaqa-based quality assessment, and automated masking of low-quality bases in error-prone assemblies from ONT reads. This standardised workflow enables reliable and scalable ONT-based genomic surveillance.

Excessive error rates in our assemblies were associated with atypical DNA modifications. Whereas elevated error rates with the SUP@v5.2 model were limited to four *S. enterica* serovar Kentucky isolates carrying a *dnd* phosphorothioation system targeting GGCC motifs, the HAC@v6.0 model generated highly inaccurate assemblies for all 13 isolates carrying either *dnd* or *dpd* systems, irrespective of the targeted sequence motif or modification type. This finding suggests that these DNA modifications were absent or underrepresented in the dataset used to train the HAC@v6.0 basecalling model. Dnd-mediated phosphorothioation and Dpd-mediated 7-deazaguanine modifications are sufficiently widespread in important bacterial pathogens that associated sequencing errors are likely to be encountered in routine genomic surveillance.

In addition, elevated error rates were observed in four *L. monocytogenes* isolates carrying methyltransferases targeting GAAGAC and GCNGC motifs. These methylation-associated errors were previously reported [18] but were substantially reduced by improved polishing with medaka v2. Nevertheless, residual motif-specific errors persisted, highlighting the continued influence of DNA modifications on nanopore sequencing accuracy. Beyond methylation, phosphorothioation, and 7-deazaguanine modifications, additional DNA modification systems likely exist and remain to be discovered. [23, 25]. Tools such as alpaqa may therefore not only support assembly quality control but also facilitate the identification of previously unrecognized modification-associated error signatures.

ONT sequencing offers several practical advantages over traditional short-read platforms, including rapid turnaround times, flexible throughput, and reduced infrastructure requirements [7]. These characteristics make ONT particularly attractive for decentralized surveillance and for resource-limited settings. However, variability in sequencing throughput between multiplexed samples remains a practical challenge. In our study, barcode imbalance was consistently observed when using the rapid barcoding kit, resulting in uneven sequencing depth across isolates (data not shown). In addition, sequencing performance varied between bacterial species; for example, *B. cereus* isolates consistently yielded lower throughput and read quality, occasionally requiring repeat sequencing. Addressing such variability will be important to ensure reliable and cost-effective implementation of ONT sequencing in routine surveillance workflows.

This study has several limitations. Although the dataset includes a large number of isolates across major foodborne pathogens, the sample size was limited for some species. Rare DNA modification systems present at low prevalence within certain species may not have been captured. In addition, Illumina and ONT sequencing were performed on independent DNA extracts. While this does not influence the systematic error patterns investigated here, it could in rare cases contribute to isolated allelic differences. Finally, our analyses were based on current nanopore chemistries and basecalling models. Future basecalling models that include more diverse DNA modification contexts may further reduce motif-associated errors identified in this study.

In conclusion, our results show that standalone ONT sequencing can provide highly accurate whole-genome assemblies suitable for routine pathogen surveillance. While rare epigenetic modifications may still introduce systematic assembly errors, these can be detected using the alpaqa computational framework, enabling correction strategies such as masking error-prone positions. Together, these findings support the transition of nanopore sequencing from a complementary technology to a standalone platform for high-resolution genomic surveillance of bacterial pathogens.

## Methods

### Isolate selection and whole genome sequencing

A total of 294 isolates representing ten major foodborne pathogens were selected for this study. The collection comprised *Bacillus cereus* group members, *Campylobacter coli*, *Campylobacter jejuni*, *Cronobacter sakazakii*, *Listeria monocytogenes*, *Salmonella enterica*, *Shigella flexneri*, *Shigella sonnei*, *Vibrio parahaemolyticus*, and *Yersinia enterocolitica* (**Supplementary Table S1**). Most isolates were collected in Switzerland and had previously been Illumina-sequenced for research or surveillance purposes at the Institute for Food Safety and Hygiene, Zurich, Switzerland. Isolates were selected based on existing Illumina data to represent a broad diversity of serovars and multi-locus sequence types (MLSTs) relevant to food safety. Within the *B. cereus* group, both cereulide-producing strains and biopesticidal *Bacillus thuringiensis* strains were included. All 78 *L. monocytogenes* isolates had been evaluated previously [18] and were included in the present study to benchmark recent bioinformatic advancements.

Genomic DNA was extracted from single purified colonies using the MagPurix Bacterial DNA Extraction Kit (Zinexts), QIAamp DNA Mini Kit (Qiagen), or DNeasy Blood and Tissue Kit (Qiagen). For short-read sequencing, libraries were prepared with the DNA Prep Library Preparation Kit (Illumina) and sequenced on the Illumina MiniSeq platform. For long-read sequencing, libraries were prepared using the SQK-RBK114 Rapid Barcoding Kit (Oxford Nanopore Technologies, ONT) according to the “Nanopore-only Microbial Isolate Sequencing Solution” protocol. Barcoded libraries were multiplexed (up to 96 isolates per run) and sequenced on MinION or PromethION devices using R10.4.1 flow cells to a minimum depth of 50x per isolate.

### Genomic analyses

Raw ONT data (pod5) were basecalled, demultiplexed, and trimmed of adapters and barcodes using dorado v1.2.0 (github.com/nanoporetech/dorado) with the SUP@v5.2, HAC@v5.2, and HAC@v6.0 models. ONT-only assemblies were generated using our Nextflow pipeline BOAP, which executes the following steps: (1) Quality filtering for a minimum read length of 1,000 bp and a minimum Q-score of 10 using nanoq v0.10.0 [26]; (2) Genome size estimation based on rapid assemblies generated with raven v1.8.3 [27]; (3) Downsampling of high-quality reads to target depths (default 100x) using filtlong 0.2.1 (parameters --length_weight 0 --window_q_weight 0) (github.com/rrwick/Filtlong); (4) De novo assembly using Flye v2.9.5 (parameter --nano-hq) [28]; (5) Contig rotation for circular sequences using DNApler v1.2.0 [29]; (6) Consensus polishing with dorado v1.2.0 using the bacterial model to produce a fastq assembly file containing per-base quality scores (-q flag) (github.com/nanoporetech/dorado); (7) Assembly contig analysis using seqkit v2.10.0 [30]; and (8) assembly accuracy assessment using alpaqa v0.1.0 (described below). For ease of installation, the BOAP version published on GitHub uses medaka v2 (github.com/nanoporetech/medaka) for consensus polishing instead of dorado polish, providing equivalent functionality.

To determine the effect of sequencing depth on assembly performance, raw reads were downsampled to predefined target depths using Rasusa v2.1.0 [31] prior to running the assembly pipeline. Illumina read adapters and low-quality bases were trimmed using fastp v0.22.0 [32]. These trimmed reads were then used to polish the ONT-only consensus assemblies with Polypolish v0.6.0 (parameter --careful) [33], and the resulting hybrid assemblies were regarded as the reference assemblies for all subsequent accuracy comparisons. To evaluate the impact of the assembly approach, selected assemblies were generated using Autocycler v0.6.1 with the automated bash script autocycler_full.sh (commit 0b72faa) [34]

Assembly completeness and the absence of contamination were confirmed using CheckM v2 1.1.0 [35]. Sequence types (STs) were identified with mlst 2.22.0 (github.com/tseemann/mlst), utilizing the PubMLST database [36]. *Salmonella* serotypes were identified *in silico* using SeqSero 1.3.1 [37]. *Bacillus cereus* group members were taxonomically assigned using Btyper3 3.4.0 [38]. Genes encoding DNA modification systems were identified in hybrid assemblies using padloc v2.0.0 [39] and abricate v1.0.0 (github.com/tseemann/abricate) (minimum coverage/identity 70%/70%) in combination with the REBASE database (downloaded on 15 April 2024) [40]. Assemblies were annotated with Bakta v1.11.1 [41]. cgMLST distances were calculated using pyMLST 2.3.0 (parameters -c 0.9 -i 0.9) [42] in combination with standardised schemes of *C. sakazakii*, *E. coli* (*Shigella*), *L. monocytogenes*, *S. enterica*, and *Y. enterocolitica* retrieved from cgmlst.org (accessed in November 2025). For *V. parahaemolyticus* and *B. cereus* sensu lato, schemes were obtained from PubMLST and modified to include only the first locus per allele for compatibility with pyMLST. During the pyMLST validation step, four problematic loci were removed from the *B. cereus* scheme. For the *B. cereus* group member *Bacillus cytotoxicus*, we developed an ad hoc scheme based on the coding sequences of the reference assembly GCF_000017425.1. To assess whether masking low-confidence bases in consensus assemblies improved the accuracy of bacterial cgMLST profiling, a subset of fastq assembly files were processed with fastq2a 0.1 (available at github.com/MBiggel/alpaqa/) to substitute bases with Phred quality scores below target thresholds with ’N’ before cgMLST analyses with pyMLST 2.3.0.

### Standalone assembly quality assessment

To identify assemblies potentially impacted by systematic basecalling errors, we developed alpaqa (“Assembly-Level Profiling And Quality Assessment”). The tool processes fastq assembly files generated with dorado polish or medaka v2 and identifies low-quality bases (LQB), defined here as bases with a Phred quality score between 1 and 5. A score of 0 was excluded to prevent bias from regions lacking quality information, such as assembly artefacts. The algorithm first masks LQB-dense regions: to avoid overcounting errors in repetitive or poorly resolved genomic regions, a sliding window (5,000 bp) scans the longest contig. Regions are masked if their LQB density exceeds five times the baseline density of the contig (minimum 0.1% density). After masking, alpaqa calculates a normalized LQB count per Megabase (Mbp) for the remaining high-quality regions. In addition, alpaqa performs a kmer enrichment analysis (k=4, 5, 6) using a binomial test to identify sequence motifs significantly associated with LQBs (*p*<0.05 after Bonferroni correction).

### LC-MS analysis of phosphorothioate (PT)-linked dinucleotides

For Liquid Chromatography Tandem Mass Spectrometry (LC-MS/MS), genomic DNA of four *S. enterica* isolates was extracted using the MasterPure Complete DNA and RNA Purification Kit (Lucigen) followed by RNAse treatment. DNA samples (1.3 to 3.0 µg) were hydrolysed with Nuclease P1 (55 °C, 2 h) and subsequently dephosphorylated using Calf Intestinal Alkaline Phosphatase (37 °C, 2 h) prior to cleanup via 10 kDa filtration. LC-MS/MS analysis was performed on an Agilent 1290 HPLC system equipped with an inline Diode Array Detector (DAD) and coupled to an Agilent 6495c triple quadrupole mass spectrometer. In each run, 10µl of hydrolysed DNA was injected onto a Phenomenex Synergi Fusion-RP column (100 × 2 mm i.d., 2.5 µm) operated at 35 °C with a flow rate of 0.350 mL/min using mobile phase solvents Buffer A (5mM ammonium acetate, pH 5.3) and Buffer B (100% acetonitrile). The gradient of buffer B was as follows: 0-15 min, 3–9%; 15-16 min, 9-95%; 16-17 min, 95-3%; 17-20 min, 3%. The DAD was operated at 260 nm to monitor signals of canonical deoxyribonucleosides. The JetStream ESI source was operated in positive-ion mode and with optimized parameters as follows: drying gas temperature, 200 °C; gas flow, 14 L/min; nebulizer, 20 psi; sheath gas temperature, 400 °C; sheath gas flow, 11 L/min; capillary voltage, 3000 V; and nozzle voltage, 1500 V. The MS was operated in dynamic multiple reaction monitoring (dMRM) mode with CE optimized for maximal sensitivity for each PT dinucleotide. Retention times of PT dinucleotides were confirmed with those from synthetic standards. Raw peak areas were extracted with Agilent Mass Hunter Workstation Qualitative Analysis software 10.0. Peak areas for each PT dinucleotide and the four canonicals acquired by DAD were converted to absolute abundances using calibration curves. The absolute abundances for PTs were then normalized by dividing by the sum of the absolute abundances of the four canonical ribonucleosides. LC-MS/MS parameters, including precursor and product ions, collision energies, and retention times for the monitored bacterial PT dinucleotides, are provided in **Supplementary Table S5**.

## Ethical statement

Ethical approval was not required because all data were collected by public health authorities as part of routine surveillance or outbreak investigations conducted under the mandate of the Swiss Epidemics Act. All patient data were anonymized.

## Acknowledgements

We are grateful to Prof. Sophia Johler for contributing Bacillus cereus isolates.

## Other notes/disclaimers

The findings and conclusions of this report are those of the authors and do not necessarily represent the official position of the Centers for Disease Control (CDC).

