## Supplements for "Standalone nanopore sequencing for foodborne pathogen surveillance: a large-scale evaluation and quality control framework"

### Supplement S1: Genetic organization of *dnd* operonss

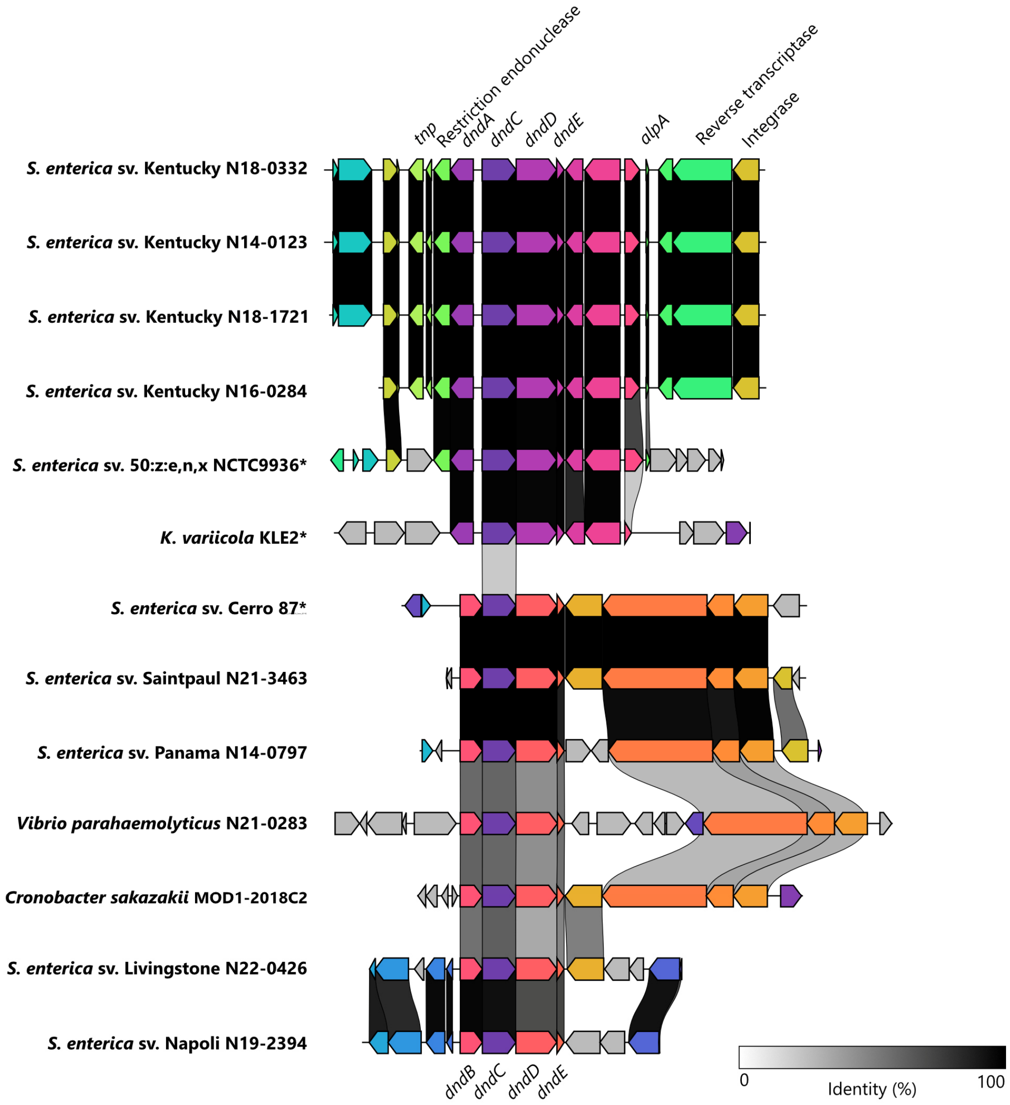


**Figure I. Comparison of the genetic structure of *dnd* operons.** The *dndACDE* operon (top), associated with *Salmonella* Kentucky, and the *dndBCDE* operon, first described *in S. enterica* serovar Cerro 87, were detected in four and six study isolates, respectively. The Kentucky-associated *dnd* operon was also identified in publicly available assemblies (in *S. enterica* serovar 50:z:e,n,x and in *Klebsiella variicola*) by BLASTn searches using the *dndC* sequence from assembly N18-0332 as a query against the NCBI core nucleotide database (Sayers et al., 2025). Shaded blocks between genomes indicate regions with ≥30% sequence identity, with grayscale intensity reflecting sequence identity as shown in the legend. Isolates marked with an asterisk (*) represent publicly available genomes (NCTC9936: GCA_900478195.1; KLE2: GCA_050990805.1; Cerro 87: GCA_001941405.1). The figure was generated using clinker (Gilchrist & Chooi, 2021).

**Supplement S2: Distribution of *dnd* in publicly available *Salmonella* serovar Kentucky assemblies**

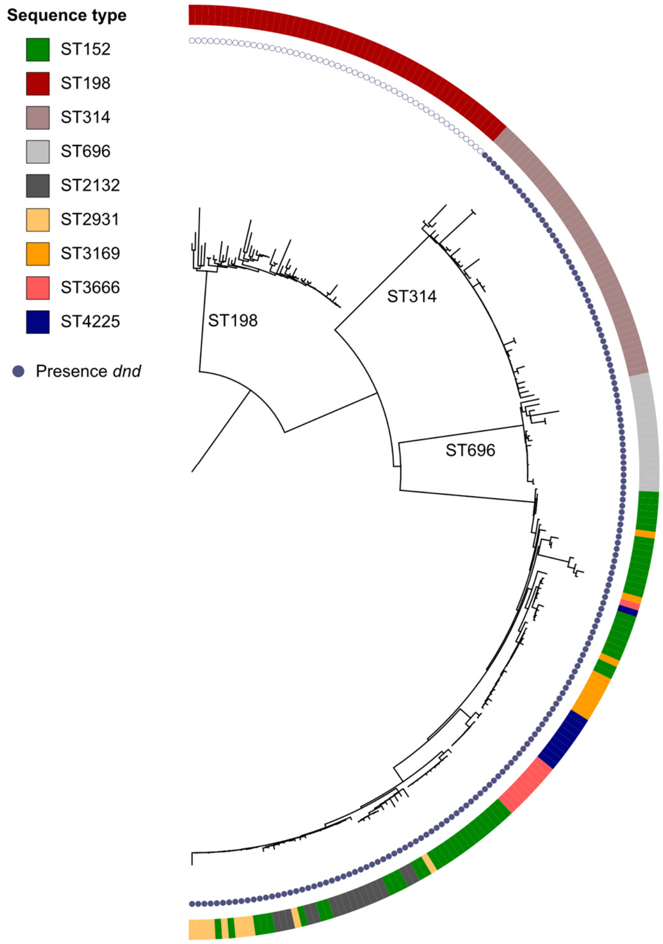


**Figure II. Phylogenetic tree of 220 *Salmonella enterica* serovar Kentucky assemblies.** Assemblies represent a subset of publicly available genomes from EnteroBase (Zhou et al., 2020) selected to capture diverse sequence types (outer ring) within this serovar. The presence of the Kentucky-associated *dnd* system is indicated by a blue filled circle. The phylogeny was estimated using Mashtree v1.4.6 (Katz et al., 2019) and visualized with iTOL v7 (Letunic & Bork, 2019).

### Supplement S3: Distribution of *dnd* and *dpd* systems in human pathogens

To determine the distribution of *dnd* and *dpd* modification systems across human pathogens, up to 5,000 genome assemblies per species were downloaded from NCBI GenBank. Assemblies were screened for antiviral defense systems using PADLOC v2.0.0 (Payne et al., 2021). A *dnd* system was considered present when at least three genes from the *dndABCDEFGH* gene cluster were detected, whereas a *dpd* system was considered present when at least nine genes from the *dpdABCDEFGHIJK* gene cluster were detected.

Overall, *dnd* modification systems were substantially more prevalent and taxonomically widespread than *dpd* systems. While 24 species lacked both systems entirely, *dnd* was detected in 44 species, with the highest prevalence observed in *Mycobacterium abscessus* (22.5%) and *Cronobacter dublinensis* (20.2%). In contrast, *dpd* systems were largely restricted to specific Enterobacteriaceae, particularly *Klebsiella* and *Escherichia* species.

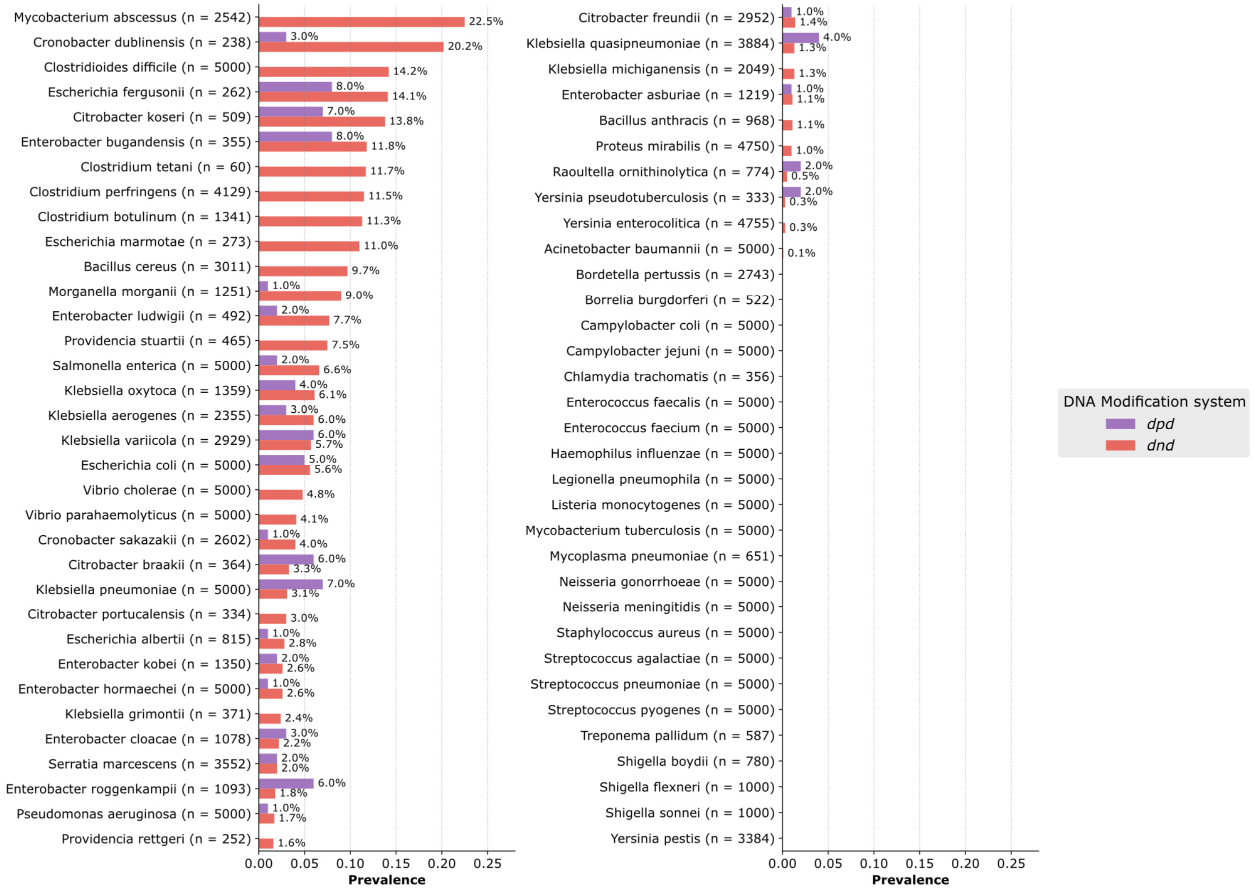


**Figure III. Distribution of *dnd* and *dpd* modification systems in bacterial pathogens.** Bar plots display the prevalence of *dnd* (red) and *dpd* (purple) DNA modification systems across 67 bacterial species (totaling 181,089 assemblies). For each species, the total number of analyzed genomic assemblies is provided in parentheses. Prevalence values (percentages) are indicated to the right of each bar.

**Supplement S4: Validation of alpaqa**

The robustness of alpaqa was evaluated using publicly available Illumina and Nanopore data from three previous benchmarking studies (Bogaerts et al., 2025; Nagy et al., 2026; Neuenschwander et al., 2026). A total of 147 isolates, mostly of clinical origin and belonging to species not included in the present study, were analyzed if they fulfilled the study criteria (minimum sequencing depth 40x for Illumina and 50x for Nanopore; SUP@v5.0 or SUP@v5.2 basecalled ONT data) (Supplementary Table S4). The dataset comprised *Escherichia coli* (n = 58), *Klebsiella pneumoniae* (n = 21), *Neisseria meningitidis* (n = 19), *Enterococcus faecium* (n = 19), *Corynebacterium diphtheriae* (n = 17), *Klebsiella oxytoca* (n = 6), *Klebsiella aerogenes* (n = 2), *Enterobacter hormaechei* (n = 2), *Citrobacter portucalensis* (n = 1), *Citrobacter freundii* (n = 1), and *Serratia marcescens* (n = 1).

Nanopore reads were randomly downsampled to a target coverage of 50x prior to assembly and analysis with the BOAP pipeline. For *Neisseria meningitidis*, the cgMLST scheme was obtained from PubMLST (Jolley et al., 2018) and modified to include only the first locus per allele for compatibility with pyMLST. For all other species, cgMLST schemes were obtained from cgmlst.org.

Of the 137 assemblies predicted to be highly reliable by alpaqa (<2.5 LQB/Mbp), 132 (96.4%) had identical cgMLST profiles compared with the Illumina-polished reference assemblies, while the remaining five showed only 1 to 3 mismatching alleles. Assemblies with higher LQB values (2.5 to 10 LQB/Mbp), predicted to be likely reliable, more frequently contained mismatches; however, all differed by at most three alleles.

**Table I.** Accuracy predicted by alpaqa and number of mismatching cgMLST alleles across 147 isolates with publicly available ONT and Illumina data.

| | LQB / Mbp | | | | |
|---|---|---|---|---|---|
| # Mismatching Alleles | **0 - 2.5** | **2.5 - 5** | **5 - 10** | **>10** | **Total** |
| **0** | 132 | 5 | 1 | 0 | **138** |
| **1 - 3** | 5 | 3 | 1 | 0 | **9** |
| **>3** | 0 | 0 | 0 | 0 | **0** |
| **Total** | **137** | **8** | **2** | **0** | **147** |

### Supplement S5: Effect of masking low-quality bases (LQBs) on cgMLST analyses

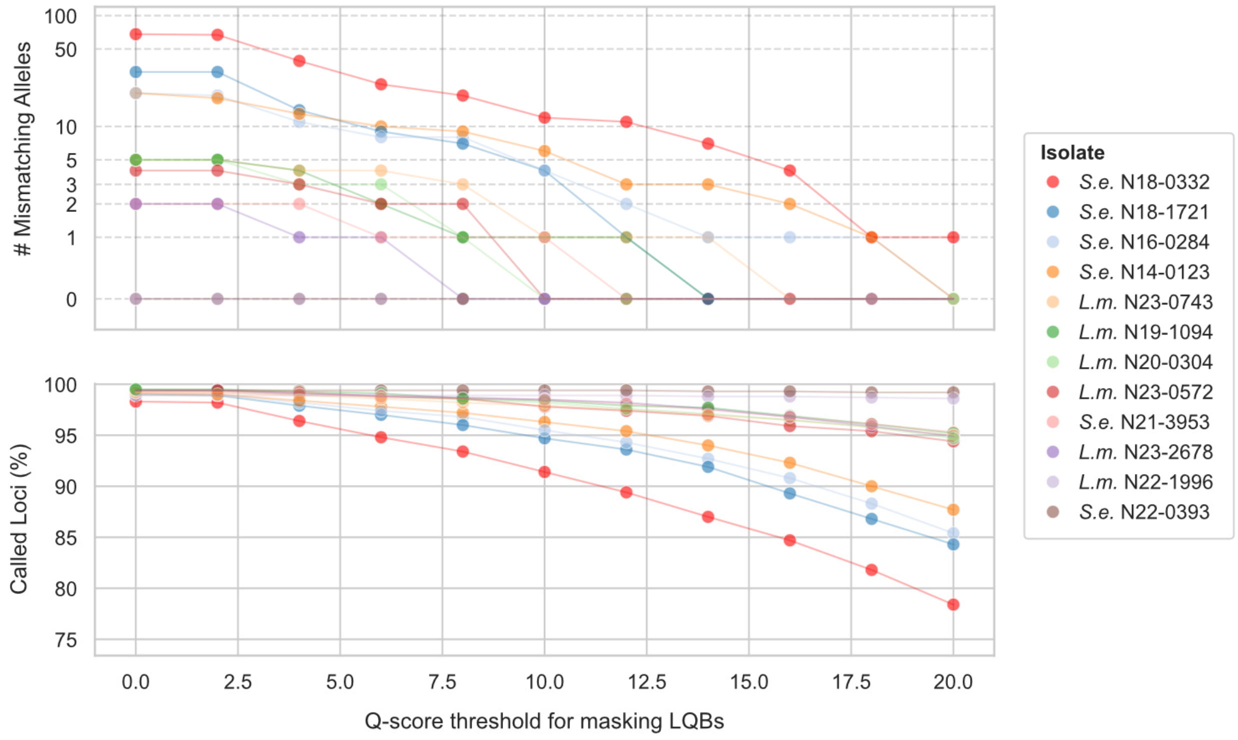


**Figure IV. Impact of masking thresholds for low-quality bases (LQBs) on cgMLST accuracy and resolution.** Top panel: Scatterplot showing the number of mismatching cgMLST alleles across different Q-score masking thresholds. Bottom panel: Scatterplot showing the percentage of cgMLST targets retained after excluding loci containing masked LQBs. The analysis was performed on six *Salmonella enterica* (*S.e.*) and six *Listeria monocytogenes* (*L.m.*) isolates, including four accurate and eight error-prone assemblies. Points are colored by isolate according to the legend. The plot illustrates that masking LQBs effectively removes unreliable alleles but also reduces the number of available targets, thereby decreasing genotyping resolution.
